# Transporting Causal Effects in Ecology: Concepts and Tools

**DOI:** 10.64898/2026.03.03.709251

**Authors:** Otto Tabell, Niklas Moser, Otso Ovaskainen, Juha Karvanen

## Abstract

1. Statistical methods related to causal inference are fundamental in ecological research as ecologists often deal with causal research questions. Consequently, recent years have seen an increase in articles discussing causal inference in ecological context. However, formally generalizing causal findings across different ecological systems still remains a challenge. While we may assess causal relationships in one location or population from experimental or observational data, replicating these findings in new settings can be impractical, expensive, or sometimes impossible.
2. We introduce causal effect transportability to ecological research — a formal framework for transferring causal effects assessed in one domain (the source) to estimate outcomes in a different domain (the target), where broader data collection may be infeasible. Using structural causal models, this framework provides formal criteria for determining when causal effects can be validly transferred between populations and derives appropriate statistical adjustment formulas when the transportation is possible. Recent algorithmic developments, implemented in accessible R software packages, automate the mathematical derivations and make transportability analysis more practical for ecologists.
3. Through simulation, we illustrate conditions under which transportability improves causal estimation relative to analyses that ignore cross-domain information. We further demonstrate the use of the methods through real-world data, examining the effect of tree canopy cover on dissolved oxygen concentrations across different watersheds.
4. Causal effect transportability offers critical tools for predicting ecological responses across heterogeneous settings, with particular relevance when experimental replication is constrained by cost, ethics, or urgency, and when management decisions require extrapolating findings to novel environmental contexts.

## 1 Introduction

Transferring causal findings across populations is a particularly relevant challenge in ecology, as ecological systems vary substantially across spatial and temporal scales (Frühstückl, 2025). For instance, experimental results from one ecosystem or region may not directly support the decision making in another ecosystem with different environmental conditions. Recent work has highlighted that ecological studies often claim “generality” without clearly defining target populations or the conditions under which findings can be transferred across ecological contexts (Spake et al., 2022). While ecologists have adopted statistical approaches to synthesize evidence across different domains, such as meta-analysis (Koricheva et al., 2013) or mixed-effects models (Bolker et al., 2009), these approaches typically treat heterogeneity as a statistical phenomenon rather than as a causal question. As a result, Spake et al. (2022) call for formal quantitative methods to assess when causal effects can be validly transferred between heterogeneous ecological systems.

Although concepts related to causal inference (Pearl, 2009) are still scarcely utilized in ecology (Franks et al., 2025), recent years have seen an increase in research articles introducing ecologists to formal causal inference (Arif and MacNeil, 2023; Byrnes and Dee, 2025; Correia et al., 2025, 2026). These articles have focused on foundational concepts such as confounding, collider bias, and mediation analysis. They demonstrate how graphical causal models can clarify long-standing ecological questions such as distinguishing direct effects from indirect effects or identifying which covariates should (and should not) be controlled for when estimating treatment effects.

Existing introductions to causal inference in ecology have largely concentrated on identifying causal effects from a single dataset. However, many ecological questions may benefit from integrating information across studies and populations rather than analyzing each dataset in isolation. Determining how such information can be combined without introducing bias is the focus of *causal data fusion* (Bareinboim and Pearl, 2016).

*Causal effect transportability* is a special case of causal data fusion that determines whether and how causal effects studied in one population (the source population) can be validly transferred for causal inference in a different population (the target population). The theoretical foundations of transportability have been formalized (Bareinboim and Pearl, 2013; Pearl and Bareinboim, 2014; Lee et al., 2020) and there are versatile computational tools (Tikka et al., 2021; Tikka and Karvanen, 2017) that can automate the derivation of transportation formulas.

In this paper, we provide one solution to the call by Spake et al. (2022) for quantitative methods that clarify when causal findings can be generalized across ecological systems. We demonstrate how the concepts and tools related to causal effect transportability can be used in practical ecological applications. We show how differences between source and target populations can be represented graphically, how the transportability of a causal effect can be assessed formally, and how the required source and target data can be combined for estimation. Finally, we illustrate the framework using simulations and an empirical application concerning the effect of tree canopy cover on dissolved oxygen across watersheds.

## 2 Causal effect transportability

We begin with a motivational example that we develop into a case study with real data in Section 3. Consider a study investigating how tree canopy cover affects dissolved oxygen concentration in stream ecosystems. Our system consists of six variables: *X* (percentage of tree canopy cover), *Y* (dissolved oxygen concentration), *W* (water temperature), *Z* (a cluster of environmental variables comprising elevation, mean precipitation and mean air temperature), *L* (land use), and *S* (season). In causal inference literature, a common way to depict the causal structure, i.e., the nonparametric causal relationships of the variables of the study, is a directed acyclic graph (DAG). An example DAG that an ecologist could draw to depict the relationships between the variables is presented in Fig. 1.

**Figure 1:**
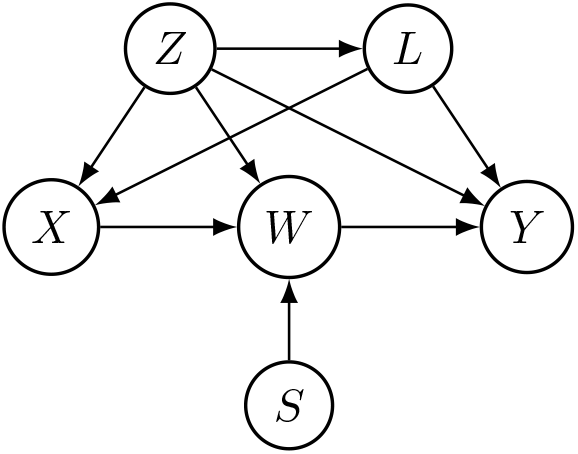
A DAG to study the effect of tree canopy cover on dissolved oxygen. Environmental variables (*Z*: precipitation, temperature, elevation) confound the effect of canopy cover (*X*) on dissolved oxygen (*Y*), while the effect of *X* on *Y* is mediated through water temperature (*W*). Unobserved land use (*L*) creates additional confounding, and season (*S*) is assumed to affect the water temperature. The environmental variables *Z* are presented as a clustered variable (Tabell et al., 2026) for brevity; they enter the statistical models in Section 3 as separate covariates. We assume canopy cover to be measured at a fixed time point to remain consistent with the real-data analysis of Section 3, and therefore the effect of season on canopy cover is not depicted in the DAG.

Suppose we are interested in knowing how dissolved oxygen (*Y*) will change in each season (*S*) if we manipulate canopy cover (*X*) to a specific level (e.g., by planting or removing trees). Assume that in geographical region A, we have observed variables *X, Y, S, Z* and *W* in a number of stream sites, but in region B, we only have data on *Z*. Land use (*L*) is not observed from either region, i.e., it is an *unobserved variable*.

Such a setting poses a fundamental question: even though we have not measured *X, Y, S, L* and *W* in region B, can we still estimate the causal effect of *X* and *S* on *Y* in this region? The answer to the question is yes: we may use causal information from region A to transport the causal effect across the two regions, as will be demonstrated later.

A naive approach that attempts to estimate the causal effect with a simple *Y* ~ *X* regression would introduce bias because *Z* and *L* confound the causal effect of *X* on *Y*. When experimental data with randomized treatment *X* is not available, we must rely on the assumed causal structure and observational data, and estimate the causal effect with methods such as back-door and front-door adjustment (Pearl, 1995). We assume that the reader is familiar with the basic concepts of causal inference, such as structural causal models (SCM) and identifiability. For readers who lack this background, we refer to existing textbooks and tutorials (Pearl, 2009; Arif and MacNeil, 2023) and provide a brief introduction to causal inference in Appendix A, expressed in the notation used throughout this paper.

Even if the causal effect can be estimated for region A, it is likely that there exist differences between the two regions, which means that generalizing the causal effect directly from region A to region B is likely to introduce bias. For instance, if the elevation, precipitation and temperature profile of region B differ from those of region A, they have, according to our causal model, a direct effect on canopy cover, water temperature and dissolved oxygen. This may, in turn, mean that the average causal effect of *X* on *Y* is different in region B.

The causal effect transportability allows us to explicitly account for such differences between the two regions. Generally, we call the domain from where we aim to transport the causal effect the *source* domain, and the domain to where we transport the causal effect the *target* domain. Causal effect transportability requires researchers to specify which aspects of the causal system may differ between the source and target domains. In a DAG, a *transportability node* pointing to a variable indicates that the causal mechanism generating that variable may differ between domains. Mechanisms associated with variables that are not children of transportability nodes are assumed to remain invariant.

An example DAG presented in Fig. 2a assumes that the marginal distributions of precipitation, temperature and elevation (i.e., *p*(*z*)) differ between the two regions, which is indicated with the transportability node *T* and the edge *T → Z*. By not including any other edges from the transportability node, we assume that any discrepancies between the regions regarding the variables *X, W* and *Y* are caused by the differences in the distribution of *Z*.

**Figure 2:**
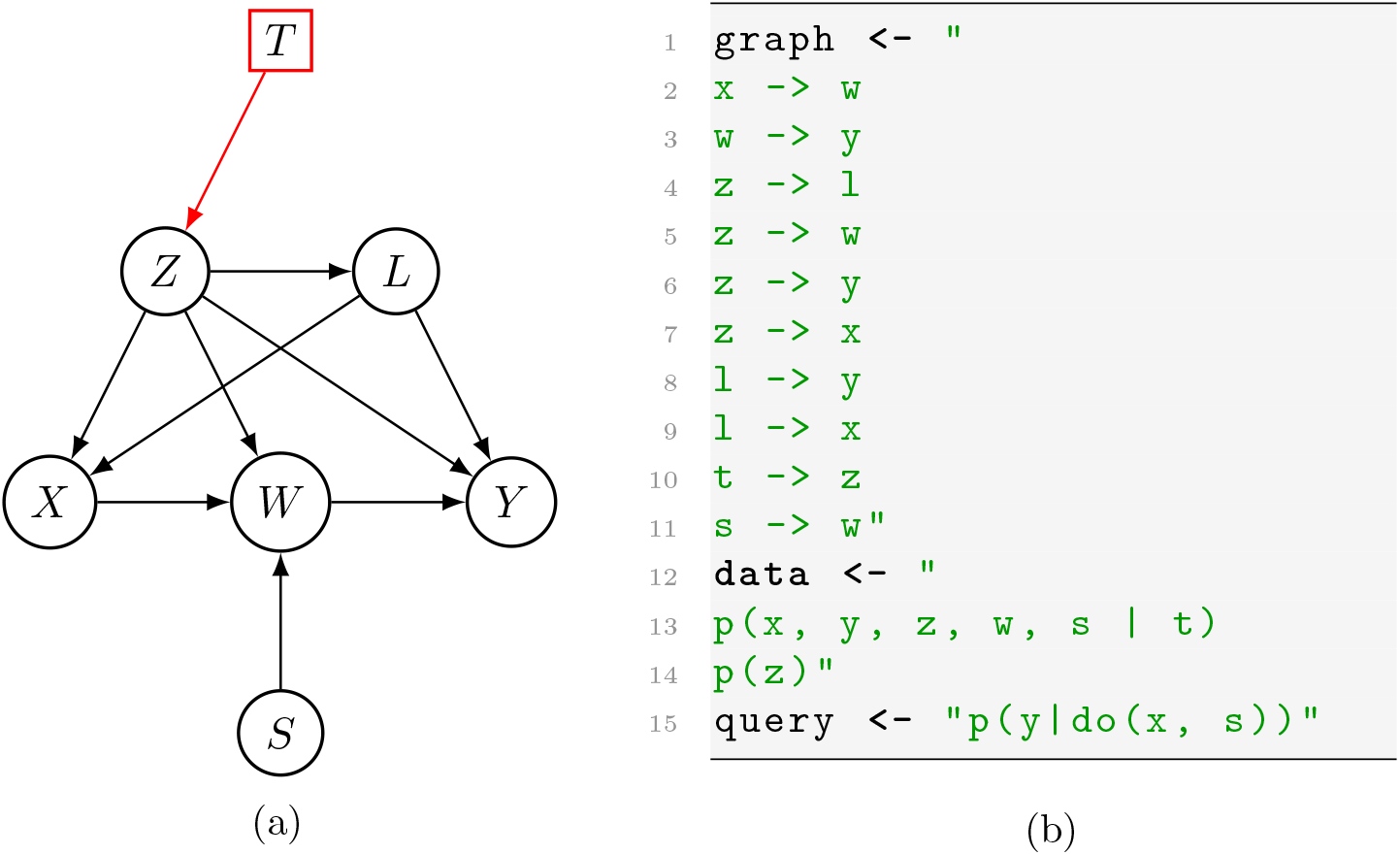
(a) The DAG of Fig. 1 augmented with a transportability node *T* → *Z*, indicating that the distribution of the *Z* (i.e., precipitation, temperature and elevation) differs between the source and target domains. (b) Defining the same DAG (lines 1–11), available data (lines 12–14) and the causal effect of interest (line 15) in text format in R.

We use notation analogous to Tikka et al. (2021), where the distributions from the source domain (Region A) are conditioned on transportability node *T* to denote the discrepancy compared to the target region. In our example, we have a random sample from distribution *p*(*y, x, w, z, s* |*t*); we use the notation *p*(*y, x, w, z, s* |*t*) to also denote the corresponding data. We assume that the observed sites constitute a representative sample of the source population and that site inclusion is not caused by variables relevant to the causal effect. Our target domain data from Region B consist only of observational measurements of the environmental covariates (*Z*). We denote this data as *p*(*z*). Note that the data are not conditioned by *T*, as it is from the target domain and therefore there does not exist a discrepancy compared to the target region.

Once the DAG augmented with the transportability node has been specified, the remaining challenge is to determine whether the causal effect of canopy cover and season on oxygen concentration, *p*(*y*|do(*x, s*)), is transportable and, if so, derive the corresponding transportation formula. This process can be automated. We next demonstrate how this can be carried out in R (R Core Team, 2025, version 4.5.1.) using the Do-search algorithm (Tikka et al., 2021) and the corresponding R package (Tikka et al., 2024).

The `dosearch`-function takes the graph, the available data sources and the query for the causal effect as inputs in a text format as demonstrated in Fig. 2b. We then call the dosearch-function that assesses the causal effect identifiability using the Do-search algorithm, given the defined graph, data and the query, while explicitly stating the transportability node:

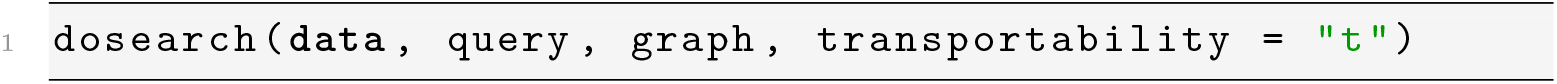

As an output, we get a text object, denoting the mathematical formulation of the identifiability formula:

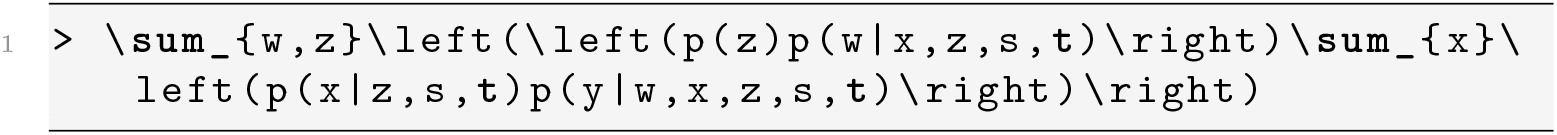

The function returns the identification formula as a LaTeX-formatted character string. When rendered in a LaTeX-compatible editor, the output corresponds to the following equation

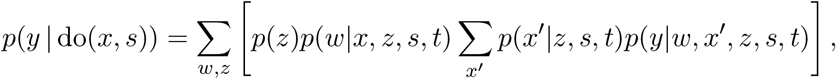

where the value of *X* is denoted by *x* in the intervention and by *x*^*′*^ in the observed data for clarity. While the equation above is valid for causal inference, we can further simplify it by noticing that *p*(*x*^*′*^ |*z, t, s*) = *p*(*x*^*′*^ |*z, t*) and *p*(*y w, x*^*′*^ |, *z, t, s*) = *p*(*y* | *w, x*^*′*^, *z, t*) because *X* and *Y* are d-separated from *S* in the corresponding distributions (see, e.g., Appendix A for further details on d-separation). Therefore, we can write

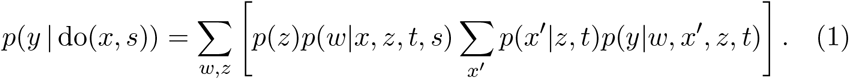

Concretely, to estimate the average causal effect from the data based on equation (1), we must follow the following steps:

1. To parametrize *p*(*w* |*x, z, t, s*), fit a model *W* ~ *X* + *Z* + *S* (Model 1) to the source population.
2. Predict values of *W* from Model 1 by plugging in the fixed *X* = *x* and *S* = *s* and *Z* from the target population.
3. To parametrize *p*(*x*^*′*^ |*z, t*), fit a model *X* ~ *Z* (Model 2) to the source population. Note the differentiation between *x* and *x*^*′*^: *x* represents the fixed value of *X*, while *x*^*′*^ represents realizations of *X* in the data.
4. Predict values of *X* from Model 2 using *Z* from the target population.
5. To parametrize *p*(*y* |*w, x*^*′*^, *z, t*), fit a model *Y* ~ *W* + *X* + *Z* (Model 3) to the source population.
6. Predict values of *Y* from Model 3 using *Z* from the target population, and the predicted values of *W* and *X* from steps 2 and 4.

We highlight that the procedure described above applies given this particular DAG and available data. If the DAG and the data differ, another procedure may be needed to estimate the causal effect. Although we have chosen linear models as Models 1–3, the identification formula itself is non-parametric. In other words, the identification formula does not assume a particular probability distribution to any element of the formula, which, in turn, means that selecting the appropriate statistical models for the ecological system under study lies in the hands of the researcher.

In addition, the procedure presented above does not itself account for the uncertainty of the estimates. To quantify the uncertainty, we suggest bootstrapping (Efron and Tibshirani, 1994) or Bayesian approaches (Gelman et al., 2013).

In some cases, the causal effect may not be transportable from the available data. For example, consider adding the edge *T → Y* to Fig. 2a. This edge indicates that the mechanism determining oxygen concentration differs between the source and target domains. Such a difference could arise, for instance, if the region B suffers from a substantially higher rate of pollution, which then alters how oxygen concentration responds to canopy cover in region B. Given this modified DAG and the available data, Do-search determines that the target-domain causal effect *p*(*y* | do(*x, s*)) is not transportable.

## 3 Case study: Portland watersheds

To illustrate the application of the methods presented in Section 2, we use observational field data on water quality from different watersheds in the greater Portland metropolitan area, Oregon, USA (Rudolph et al., 2025). These data are integrated with data on the landscape characteristics (Hopkins et al., 2024) of the sampling sites. Our task is to estimate how an intervention on tree canopy cover (e.g., cutting or planting trees) affects oxygen concentration. We consider four hypothetical canopy-cover interventions, that set the canopy cover to 45%, 50%, 55%, or 60%, and study their effect together with the effect of the season.

To illustrate causal effect transportability, we assume that oxygen measurements are unavailable from one watershed (Fanno Creek), which serves as our target domain — a scenario realistic in ecological monitoring where, for instance, resource constraints can limit data collection to a subset of sites. Our aim is to transport the causal effect estimated from other watersheds grouped together (our source domain) to estimate oxygen responses to canopy cover in Fanno Creek.

We first evaluate the framework in a controlled simulation setting where the true causal effect is known. The simulation uses the real Portland data as its basis. We then show how the same methods can be applied to real data, and discuss their benefits and potential limitations.

### 3.1 Data

The data were collected across four seasons between summer 2023 and spring 2024, with all samples within each season collected within a five-day window. Several physicochemical quantities were measured from the water samples. In this case study, we are using the dissolved oxygen concentration (mg/L) and water temperature (°C). The landscape of the sampling sites is characterized by the tree canopy cover (TCC) percentage, the mean elevation (meters above the sea level), mean annual precipitation 1991–2020 (mm) and the mean temperature 1991–2020 (°C). TCC has been estimated from satellite images from 2021. Consequently, the variables describing TCC, the mean precipitation, the mean temperature and the mean elevation stay constant through different dates in our data. This also explains our choice to leave the edge *S → X* out of the DAG in Fig. 2a.

Prior to analysis, several data-cleaning steps were applied. The Columbia Slough watershed was excluded from the dataset, as its characteristics differed substantially from the other watersheds in ways not captured by the covariates in our statistical models. One duplicate spring-season measurement recorded for a site in the Willamette watershed was removed, as was one site lacking a tree canopy cover estimate. Finally, samples with an implausible dissolved oxygen percent saturation, indicative of measurement error, were excluded. After these exclusions, the source population (all watersheds grouped excluding Fanno Creek) comprised *N* = 247 observations, and the target population (Fanno Creek) comprised *N* = 100 observations.

The locations of the sampling sites are illustrated in Fig. 6 of Appendix C. The locations of the Fanno Creek sampling sites are clearly geographically distinct from the rest of the sites. The median oxygen concentration in Fanno Creek (9.31 mg/L) differs considerably from the median oxygen concentration in the rest of the watersheds (10.11 mg/L), further suggesting that systematic differences between the target and source populations exist that may threaten direct generalization. We have illustrated the relationships between the canopy cover and oxygen concentration per watershed and season in Fig. 7 of Appendix C. In this case study, we assume that the difference is caused by functional discrepancies between the target and source in the environmental background variables elevation, temperature and precipitation. These differences are depicted in Fig. 8 of Appendix C.

### 3.2 Methods and models

We assume the DAG of Fig. 2a. For the purpose of simplicity, to estimate *p*(*y*| do(*x, s*)) based on equation (1), we use linear models with Gaussian error terms in the case of the three models required to be fit during the procedure. As a large majority of TCC values in Fanno Creek fall between 45 and 60% (see, e.g., Fig. 7), we estimate the dissolved oxygen concentration for four fixed TCC values of 45, 50, 55 and 60.

In this case study, we use non-parametric bootstrapping to quantify the uncertainty. The source (N = 247) and the target data (N = 100) are both resampled with replacement, to generate new bootstrap source and target data. Using the new source and target samples, we estimate the causal effect *p*(*y*|do(*x, s*)) using the six-step procedure presented earlier. This process is repeated 2 500 times to obtain the distribution for the average causal effect.

We compare three modeling approaches: fitting and predicting only in the source population, i.e., a model that does not take the target population into account (Source population model); b) transportation by fitting the model in the source population but predicting in the target population using the six-step procedure described in Section 2 (Transported model); c) fitting and predicting only in the target population which also serves as the kind of model we would fit should we have full data available for Fanno Creek (Target population model). The Source and the Target population models also use the six-step procedure, but for the Source population model all data used comes from the source population while for the Target population model all data comes from the target population.

### 3.3 Simulation

For the simulation, we generate data using the Portland case study as the basis. We assume the causal structure of Fig. 2a, and use the observed values from the data for the landscape variables *Z* and season *S*, and then generate variables *L, X, W* and *Y* as explained in Appendix B. For instance, *X* is defined as a function of *Z* and *L* and random normally distributed noise, as is depicted in Fig. 2a.

We compare the performances of the three different models that were presented earlier. In a controlled setting, if the transportation is successful, the results by Transported and Target population models should resemble each other, while estimates from the Source population model should have some bias as they do not take the target population into account at all. We fix canopy cover to 55% and aim to estimate the dissolved oxygen concentration in the target population with the fixed canopy cover for each season. We generate a total of 2 500 simulation instances.

### 3.4 Results

The results for the simulation study are presented in Fig. 3. The Source population model overestimates the average oxygen values for each season, which signals that we cannot naively generalize the causal conclusions based on a prediction model fitted to the source data. As expected, estimates by the Transported model closely matched the true values. There is a minimal difference compared to the Target population model.

**Figure 3:**
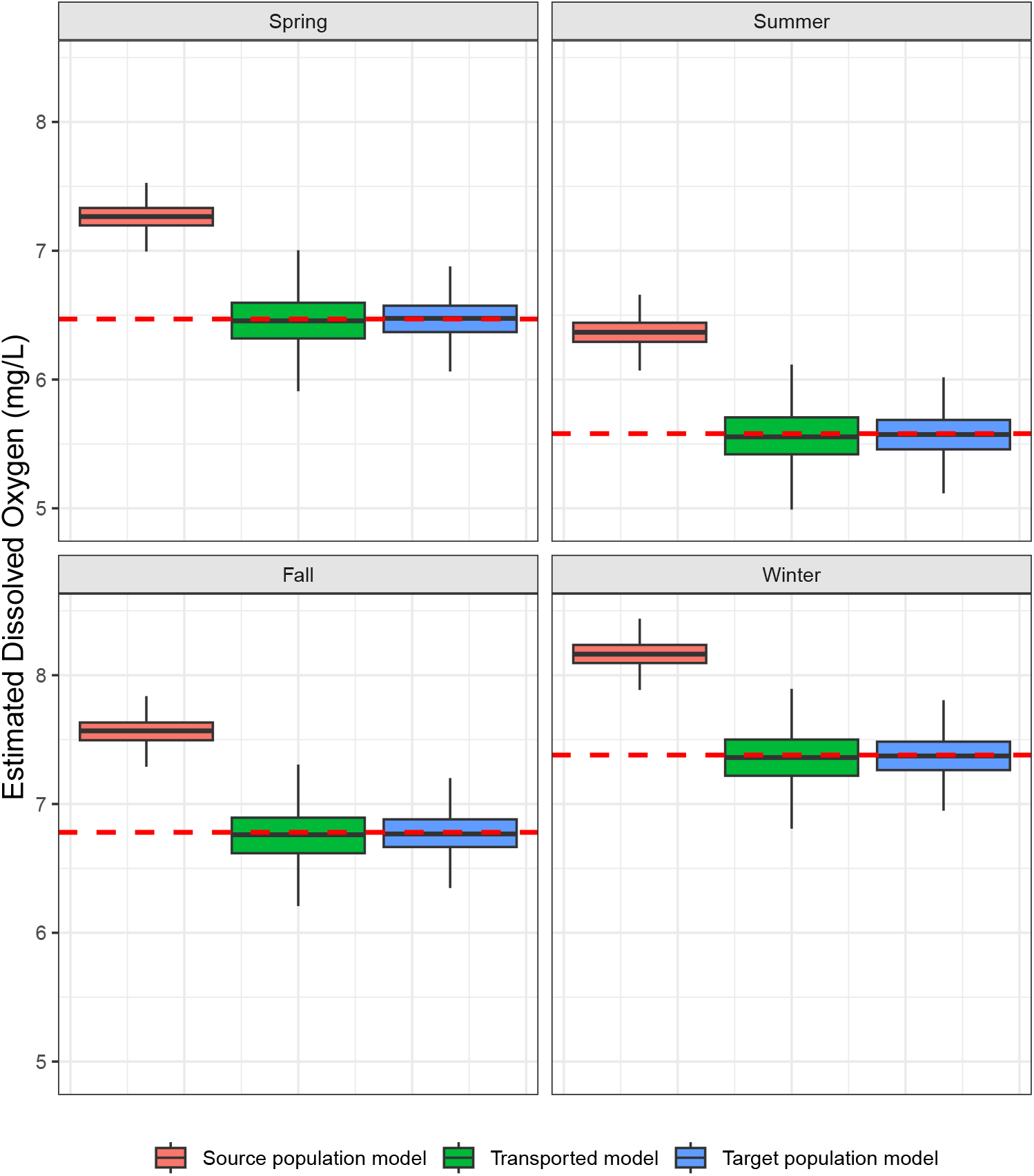
The results for the simulation study. The dissolved oxygen concentrations are reported separately for each season under the intervention do(*X* = 55%), using three modeling approaches. Boxplots show median, interquartile range (IQR), and whiskers extending to 1.5 *×* IQR calculated from 2500 simulation runs; outliers beyond the whiskers are omitted for clarity. The red horizontal lines represent the true dissolved oxygen concentration values: 6.47, 5.58, 6.78 and 7.38 mg/L for spring, summer, fall and winter, respectively.

The results for the real data are presented in Fig. 4. In the real-data analysis, the average transported estimates generally move from the source-only estimates towards the target-only estimates. However, unlike in the simulation study, the transported estimates differ from the target-only model. As the ground-truth is unknown, these results do not measure the performance of the models but act as an illustration of transportation with real ecological data. The real processes may differ from our assumptions in many ways. For instance, the DAG may miss some confounders, the domains may differ also with respect to other variables than *Z*, and the linear models chosen because of their simplicity might be inadequate for true functional relationships.

**Figure 4:**
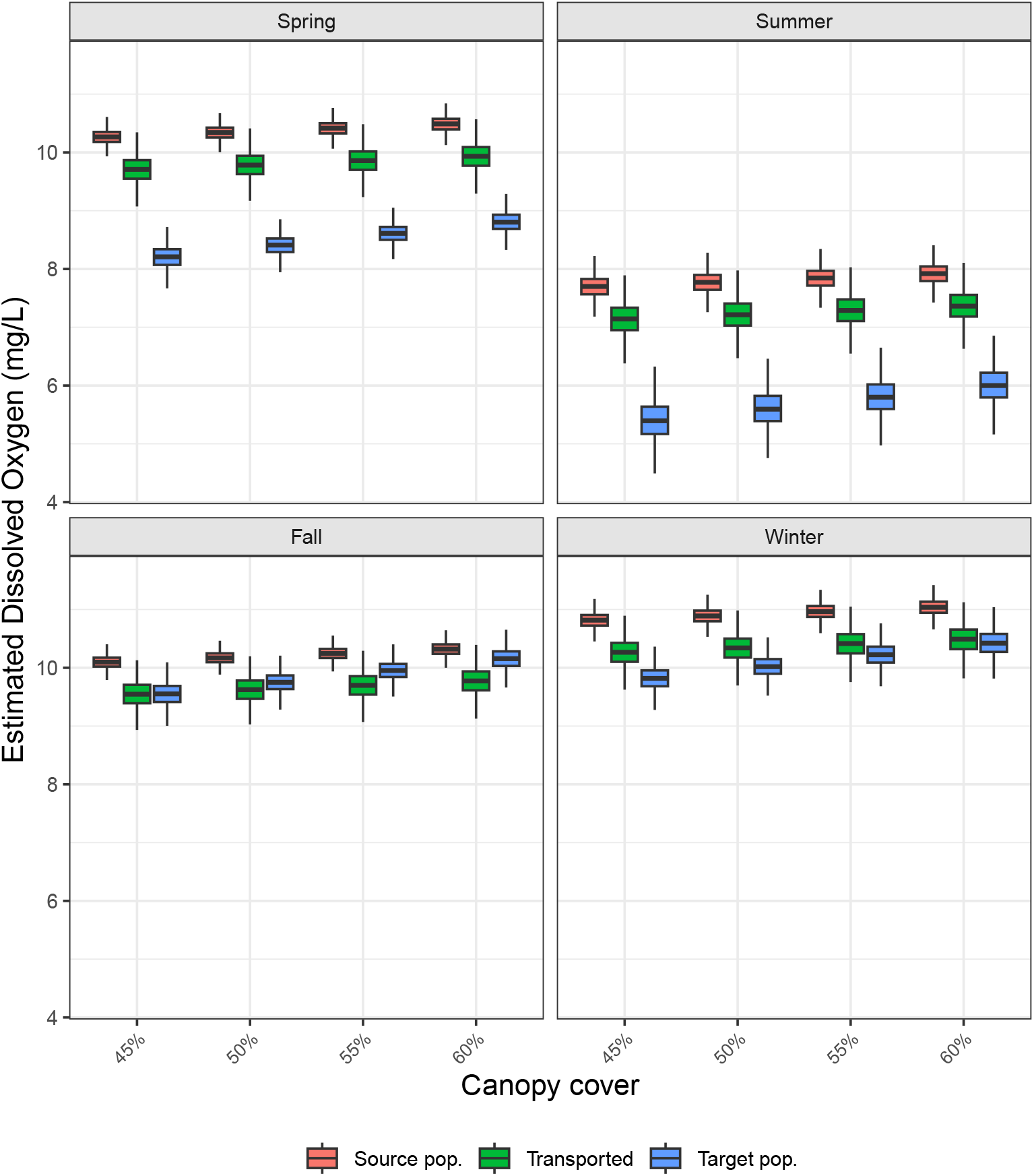
The results for the real-data case study. The dissolved oxygen concentrations are reported separately for each season under the interventions do(*X* = 45%), do(*X* = 50%), do(*X* = 55%) and do(*X* = 60%), using three modeling approaches. Boxplots show median, interquartile range (IQR), and whiskers extending to 1.5 *×* IQR calculated from 2500 bootstrap iterations; outliers beyond the whiskers are omitted for clarity.

Additionally, we replicated the analyses of the case study for two other watersheds (Johnson Creek and Kellogg Creek) for which we had at least 10 samples available for each season. The analysis results are presented in Figs. 9 and 10 of Appendix C. The general conclusions are similar to those from the Fanno Creek case.

## 4 Discussion

In this paper, we demonstrated that transporting causal effects across heterogeneous ecological settings can serve as a useful tool in ecological research. Our proposed methods could particularly be beneficial when collecting data from a new domain is constrained by cost, ethics or urgency, for example. The use of the methods was illustrated in the case study using real urban stream data from Portland, Oregon. The related simulation study showed that a naive approach that does not take the underlying causal structure into account, can lead to biased causal conclusions.

Transporting causal effects is highly relevant, for instance, under the European Union’s Water Framework Directive which obliges its member countries to monitor the status of the surface and groundwater, and intervene in the management if necessary (European Parliament, 2000). As ecological monitoring can be financially costly in some cases (e.g., Koski et al., 2020), transportability methods could enable managers, for instance, to prioritize monitoring investment by predicting which locations are most likely to benefit from interventions, extend causal findings from intensively studied reference sites to less-monitored locations, or forecast intervention outcomes before committing resources to implementation and monitoring.

In causal inference, fundamentally incorrect causal assumptions may lead to misleading conclusions. This also extends to causal effect transportability: if key causal pathways are omitted from the DAG, important transportability functional discrepancies are not represented in the graph, or causal mechanisms genuinely differ across domains, transported estimates may be biased despite satisfying the identification formula. In addition, while we demonstrated that the causal effect identification can be carried out by algorithms, we highlight that the whole process is not merely mechanical. Domain knowledge is still required in deriving the DAG, data acquisition, and transferring the identifiability formulas to statistical models.

In this paper, we focused on simple examples with two domains and data sources. Causal effect transportability can be extended to settings with more than two domains and sources (see, for example, Lee et al. (2020)), and Do-search can be also used in such cases. The framework can also be applied, for instance, to integrate more flexible modeling solutions such as generalized additive models (Hastie and Tibshirani, 1986) or spatio-temporal modeling (Cressie and Wikle, 2011) that could address non-linearity and spatial and temporal autocorrelation — issues typically arising in ecological data. In addition, other cases of applying causal data fusion, such as recovering from selection bias (Bareinboim and Pearl, 2012) or dealing with multiple partially overlapping data sources (Tikka et al., 2021), could be well applied also to ecological research problems.

To conclude, we believe that the proposed framework responds directly to the call by Spake et al. (2022) for quantitative methods to assess when and how causal effects can be validly transferred between populations. More broadly, the structural causal modeling framework provides ecologists with rigorous means to identify causal effects from observational data, transport those effects to domains where direct data collection is constrained, and integrate experimental and observational data sources that alone would be insufficient to answer the causal question of interest.

## Author contributions

Otto Tabell, Niklas Moser, Otso Ovaskainen and Juha Karvanen contributed to the conceptualization and methodological framing of the manuscript, and to reviewing and editing the paper. Otto Tabell and Niklas Moser reviewed the literature. Otto Tabell led the writing of the manuscript and conducted the data-analyses and simulations.

## Acknowledgments

The authors wish to thank the anonymous referees that helped to improve the paper. Claude Sonnet 5 was used during revision to draft and edit portions of figure codes in Appendix C. All output was verified and edited by the authors, who take full responsibility for the content. This work was supported by the Finnish Doctoral Program Network in Artificial Intelligence (AI-DOC), Decision VN/3137/2024-OKM-6. Otso Ovaskainen and Niklas Moser were funded by the Research Council of Finland (grants no. 336212 and 345110 to OO). Juha Karvanen was supported by the Research Council of Finland (grant no. 368935).

## Data availability statement

The water sample data from Portland, Oregon is available at https://doi.org/10.6073/pasta/5ce0fb531349f8be9e9d5b1ae6d7e59a (Rudolph et al., 2025). The landscape data for the corresponding sites is available at https://doi.org/10.5066/P13UZYZF (Hopkins et al., 2024). The R codes used in the case study and for deriving the identifiability formulas in `dosearch` are available at https://github.com/ottotabell/transportability-ecology.

## A Brief introduction to structural causal models

In the paper, we assume the reader to be familiar with the basic concepts of structural causal models. While we acknowledge the wide range of methods for causal inference from observational data (Imbens and Rubin, 2015; VanderWeele, 2015; Tchetgen Tchetgen et al., 2024; Grace, 2021), we highlight that the structural causal model framework offers state-of-the-art tools particularly to address the causal effect transportability (Bareinboim and Pearl, 2013; Lee et al., 2020; Tikka et al., 2021), which is why it is the chosen toolkit in the paper. In this appendix, we briefly introduce the interested readers to the basic concepts of structural causal models.

Here, we consider a study investigating how grazing affects plant richness in grassland ecosystems. Our system consists of six variables: *X* (grazing intensity), *Y* (plant species richness), *W*_1_ (vegetation height), *W*_2_ (light availability at ground level), *Z*_1_ (soil moisture), and *Z*_2_ (slope/topography). An example DAG that an ecologist could draw to depict the relationships between the variables is presented in Fig. 5.

**Figure 5:**
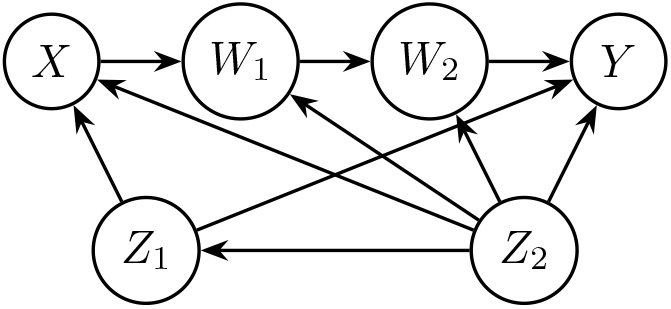
DAG for the grassland grazing system example. We assume that grazing intensity (*X*) affects plant species richness (*Y*) directly and indirectly by reducing vegetation height (*W*_1_), which increases light availability at ground level (*W*_2_). Soil moisture (*Z*_1_) influences where herbivores preferentially graze, which plant species can establish, and vegetation growth. Slope (*Z*_2_) is assumed to affect all aforementioned variables. For illustrative purposes, some other possible causal relationships, such as *Z*_1_ → *W*_1_ or the direct effect *X* → *Y*, are not included in the graph.

### A.1 Graph theoretical concepts and d-separation

A *path* is a sequence of nodes connected by arrows where each variable is linked to the next one, regardless of arrow direction. For example, in Fig. 5, there exist several paths between *X* and *W*_2_, such as *X ← Z*_2_ → *W*_2_. A *directed path* from a node *A* to a node *B* is a directed path in which every arrow points away from *A* and towards *B*. For example, in Fig. 5 the only directed path from *X* to *W*_2_ is *X → W*_1_ → *W*_2_. If a directed path from *A* to *B* exists, *B* is a *descendant* of *A*. A *cycle* is a path that starts and ends at the same variable. Fig. 5 does not contain cycles and is therefore *acyclic*. A path between two nodes, say *A* and *B*, is *d-separated* by a variable set **S** if

- the path contains a *chain C → M → D* or a *fork C ← M ←D* such that *M* is a member of **S**.
- the path contains a *collider C → M ← D* such that neither *M* nor any of its descendants belong in **S**.

Variables *A* and *B* are d-separated by **S** if **S** d-separates every path between *A* and *B*.

For instance, in Fig. 5 the set **S** = {*Z*_2_, *W*_1_} d-separates *X* and *W*_2_: *Z*_2_ blocks the fork paths *X ← Z*_2_ → *W*_2_ *X ← Z*_2_ → *W*_1_ → *W*_2_, while *W*_1_ blocks the chain *X → W*_1_ → *W*_2_. Additionally, any paths through *Y* are blocked because it acts as a collider and it is not included in the set **S**. We denote the d-separation of *X* and *W*_2_ given *S* in a DAG *G* as (*X* ╨ *W*_2_ | *S*)_*G*_. Additionally, we denote a graph *G*, where all incoming arrows to *X* are removed as 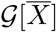, and a graph where all outgoing arrows from *X* are removed as *G*[*<u>X</u>*].

### A.2 The do-operator and the interventional distribution

A causal research question concerns how an outcome would behave under an intervention, rather than how the outcome is associated with an observed value of another variable. This distinction is represented using the do-operator. For example, *p*(*y* | do(*X* = *x*)) denotes the distribution of the outcome *Y* under a hypothetical intervention that sets the treatment *X* to the value *x*. The intervention do(*X* = *x*) replaces the process that normally determines *X* with an external assignment of the value *x*.

In a DAG, this intervention is represented by removing all arrows entering *X*, while retaining arrows leaving *X*. The resulting graph represents a system in which *X* no longer depends on its usual causes. This graphical operation formalizes an idealized intervention, such as the random assignment of a treatment. It does not, by itself, provide a numerical estimate of the interventional distribution.

The interventional distribution *p*(*y* | do(*X* = *x*)) should be distinguished from the observational conditional distribution *p*(*y* | *X* = *x*). The latter describes the distribution of *Y* among observational units for which *X* = *x*. Consequently, *p*(*y* | do(*X* = *x*)) ≠ *p*(*y X* = *x*) in general. The two distributions may coincide under particular causal assumptions, for example when there is no confounding between *X* and *Y*.

### A.3 Back-door adjustment and average causal effect

A widely-used way to address confounding is the *back-door adjustment* (Pearl, 1995) where the aim is to condition the response on an *admissible set*. An admissible set is a set of variables that d-separates all paths between the treatment and the response that end with an arrow to the treatment. Such a path is called a *back-door path*. In addition, this set may not contain variables that are descendants of the treatment, i.e. nodes to which there is a directed path from the treatment variable.

In the case of Fig. 5, there exist several back-door paths between *X* and *Y*, including but not limited to *X ← Z*_1_ → *Y* or *X ← Z*_2_ → *W*_1_ → *W*_2_ → *Y*. We can notice that *Z*_1_ and *Z*_2_ are variables that block all such back-door paths, and therefore we can consider {*Z*_1_, *Z*_2_} as our admissible set. This set may not include *W*_1_ or *W*_2_ because they are descendants of *X*.

Using the back-door adjustment, we can estimate the causal effect *p*(*y* | do(*x*)) by adjusting with the admissible set. In other words, we have

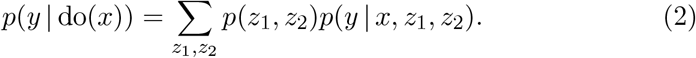

The distribution *p*(*y* do(*x*)) can be summarized by its expectation E[*Y* do(*X* = *x*)] which is called *average causal effect*. Especially for binary treatments, average causal effect may also refer to difference E[*Y* | do(*X* = *x*_1_)] −E[*Y* | do(*X* = *x*_0_)]. Typically, we are interested in translating the identification formula into concrete statistical models in order to estimate the average causal effect.

In practice, this is essentially what we do when “controlling for” covariates in regression models. We would fit a model predicting species richness (*Y*) by grazing intensity (*X*), soil moisture (*Z*_1_) and slope (*Z*_2_), and then use this model to predict richness for different grazing intensities, while averaging over their observed distributions. This adjustment removes the confounding bias introduced by *Z*_1_ and *Z*_2_. Importantly, we do not adjust for *W*_1_ or *W*_2_ because they are mediators on the causal path from grazing to richness. Adjusting for them would block the causal path we want to estimate.

The back-door adjustment formula of equation (2), where *X* receives a fixed value *x*_fix_, can be written as

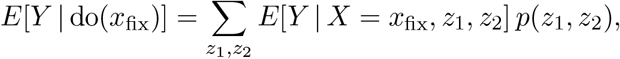

i.e., we are weighting the conditional expected value *E*[*Y* | *X* = *x*_fix_, *z*_1_, *z*_2_] by the distribution *p*(*z*_1_, *z*_2_).

We therefore need an estimator *Ê* [*Y X* = *x*_fix_, *z*_1_, *z*_2_], a statistical model where we predict *Y* by *X, Z*_1_ and *Z*_2_. Assume that *X, Z*_1_ and *Z*_2_ have independent linear effects on *Y* and that the errors of *Y* are normally distributed. Using a sample of *n* observations, we can then fit a linear model

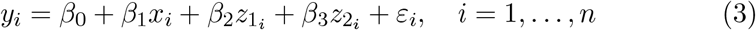

where *β*_0_, *β*_1_, *β*_2_ and *β*_3_ are regression coefficients to be estimated and *ε*_*i*_ ~ *N* (0, *σ*^2^)

The weighting by *p*(*z*_1_, *z*_2_) can be conducted by averaging *Ê* [*Y* | *X* = *x*_fix_, *z*_1_, *z*_2_] over the observed values of *Z*_1_ and *Z*_2_. In other words, we have

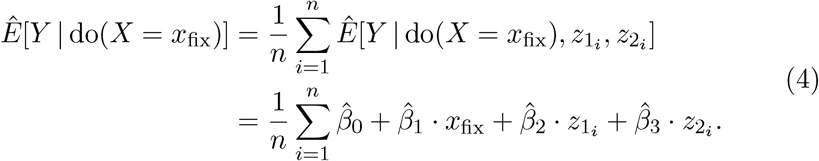

In our grazing-richness example, equation (3) gives an estimate on how species richness responds to grazing intensity, soil moisture and slope. To estimate the average causal effect of setting grazing to a specific intensity *x*_fix_, the equation (4) averages the predicted values over the observed range of slope and soil moisture values. This effectively asks: “What would richness be at each observed moisture and slope level if we experimentally set grazing to *x*_fix_?”

### A.4 Do-calculus

Identification of causal effects relies on *do-calculus* (Pearl, 1995), which consists of three rules for manipulating interventional distributions:

1. Insertion and deletion of observations:

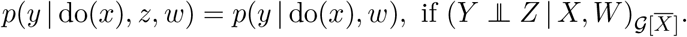
2. Exchanging actions and observations:

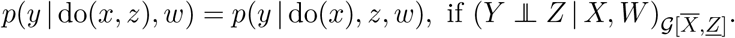
3. Insertion and deletion of actions:

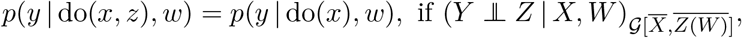

where *Z*(*W*) denotes variables in the set *Z* that do not contain a directed path to the set *W*.

Consider the graph of Figure 5 with the aim of identifying the causal effect *p*(*y*| do(*x*)) and assume all variables are observed. Earlier, we already concluded that the causal effect can be identified with back-door adjustment using the equation (2). We can also derive the same formula by do-calculus and standard probability manipulations accordingly. First, marginalize over *Z*_1_ and *Z*_2_ and use the chain rule:

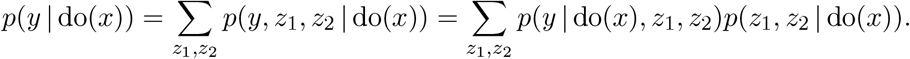

Because (*Y* ╨ *X* | *Z*_1_, *Z*_2_)_*G*[*X*]_, we can use rule 2 of do-calculus to have:

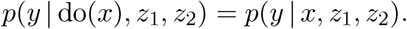

Additionally, because (*Z*_1_, *Z*_2_ ╨ *X*), we can use rule 3 of do-calculus for:

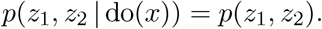

Combining the results above, we can write:

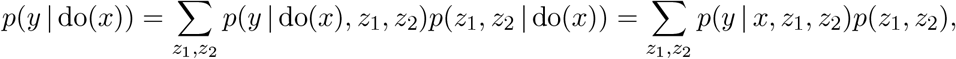

corresponding to the back-door criterion.

### A.5 Notations for the data sources

The variables in a DAG can be classified as observed, intervened, conditioned or unobserved. In Table 1, we summarize different types of data sources, and their corresponding notations based on the classification.

**Table 1:** Four types of data sources considered in this paper, with example notations and brief explanations. The variables are related to the DAG of Fig. 5. In each source of data, the variable *W*_2_ is considered *unobserved*.

| Data source | Example notation | Brief explanation |
| --- | --- | --- |
| Observational | $p(x, y, w_1, z_1, z_2)$ | Data are collected on $X$, $Y$, $W_1$, $Z_1$, $Z_2$ without intervention or conditioning. The full joint distribution is observed. |
| Experimental | $p(y, w_1, z_1, z_2 \mid \text{do}(x))$ | The variable $X$ is externally intervened upon, and observations of $Y$, $W_1$, $Z_1$ and $Z_2$ are drawn from the resulting interventional distribution. |
| Conditional | $p(x, y, w_1 \mid z_1, z_2)$ | Data on $X$, $Y$ and $W_1$ are collected only for units satisfying particular values of $Z_1$ and $Z_2$. |
| Experimental & conditional | $p(y, w_1, z_1 \mid \text{do}(x), z_2)$ | Data are obtained under an intervention on $X$, while the sampling is simultaneously restricted to units with a specified value of $Z_2$. |

## B Simulated data

In this section, we explain how the data used in the simulation study of Section 3 were generated. We begin the construction of the simulated data by using the values in the real data for variable *Z*, i.e. precipitation, temperature and elevation.

Land use is assigned to each site using a multinomial logistic model. For watershed *i*, we define the linear predictors for each land use category:

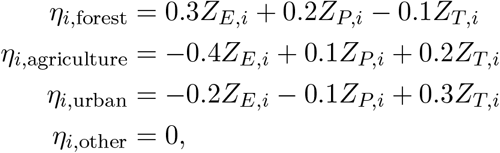

where *Z*_*P,i*_, *Z*_*T,i*_ and *Z*_*E,i*_ are standardized precipitation, temperature and elevation, respectively.

The unnormalized probabilities are computed using the exponential function:

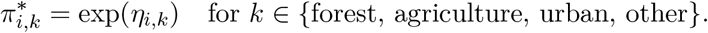

These are then normalized to ensure they sum to 1:

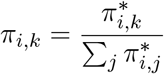

Finally, land use for site *i* is sampled from a categorical distribution:

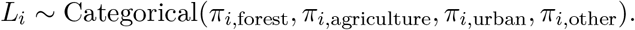

Tree canopy cover is generated as a function of both land use type and environmental conditions:

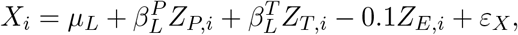

where *ε*_*X*_ ~ *N* (0, 4^2^) and the land use-specific parameters are:

- Forest: *µ*_forest_ = 65, 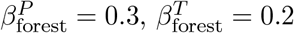
- Agriculture: *µ*_ag_ = 50, 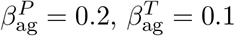
- Urban: *µ*_urban_ = 55, 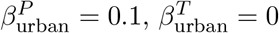
- Other: *µ*_other_ = 58, 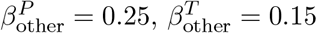

In addition, we constrain the canopy cover to the range [45%, 75%].

Water temperature (W_*is*_) is modeled with seasonal baselines (BL_*s*_) and environmental drivers:

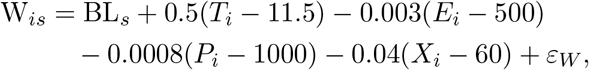

where *s* indexes seasons with baselines (*BL*_*s*_): Spring (12*°*C), Summer (15*°*C), Fall (11*°*C), Winter (9*°*C). The error term *ε*_*W*_ ~ *N* (0, 1.0^2^).

The outcome variable, dissolved oxygen, is influenced by land-use, environmental variables and water temperature:

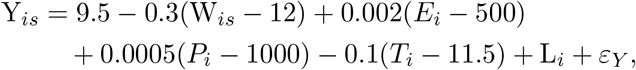

where land use effects are: Urban (−1.2 mg/L), Agriculture (−0.6 mg/L), Forest (+0.8 mg/L), Other (0 mg/L), and *ε*_*Y*_ ~ *N* (0, 0.4^2^).

## C Additional analyses in the case study

This appendix provides additional analyses regarding the case study. We first provide a map illustrating the locations of the sampling sites in Fig. 6. We then provide illustrative figures regarding the variables of the study in Figs. 7 and 8. We then selected two additional watersheds (Johnson Creek and Kellogg Creek) that had at least 10 observations per season, in order to replicate the transportability analyses performed in Section 3. For Johnson Creek, we used the same fixed canopy cover values (i.e., 45, 50, 55 and 60%), while for Kellogg Creek we used canopy cover values shifted to 35, 40, 45 and 50%, as this watershed had lower canopy cover percentage compared to the other watersheds. The results for the two other watersheds are presented in Figs. 9 and 10.

**Figure 6:**
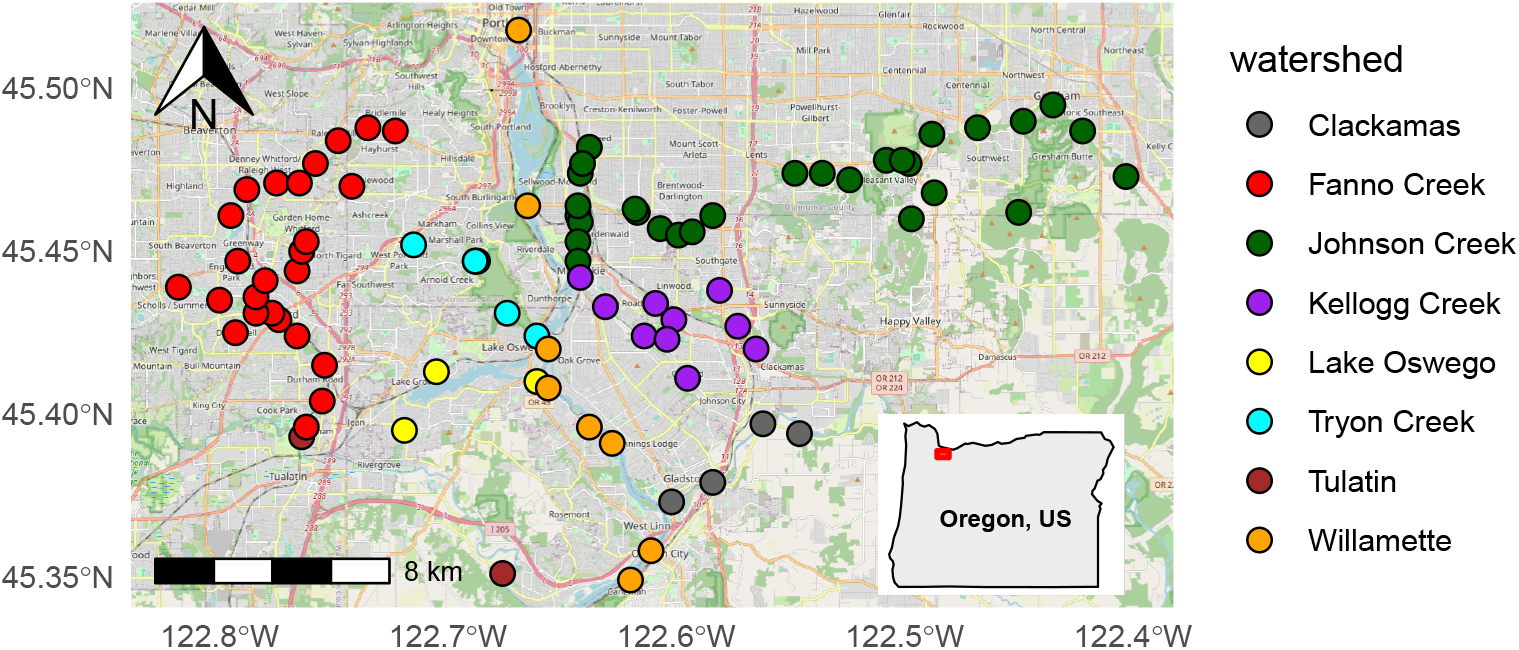
A map illustrating the locations of the sampling sites of the case study. The points are colored by the watershed. Base map data © OpenStreetMap contributors, available under the Open Database License (https://www.openstreetmap.org).

**Figure 7:**
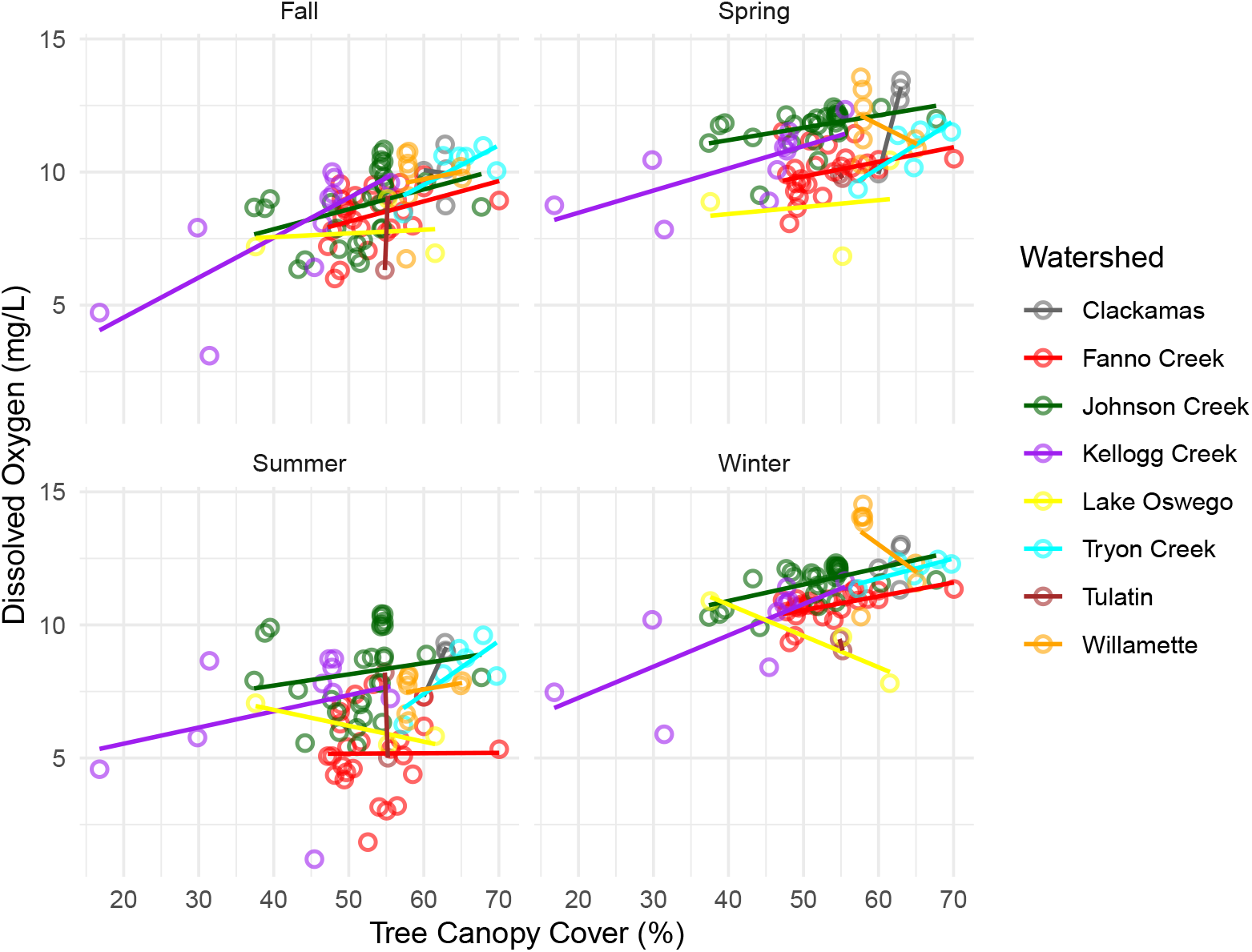
Regression lines between tree canopy cover and dissolved oxygen concentrations across Portland watersheds per season. Each point represents a single water sample, with colors indicating watersheds.

**Figure 8:**
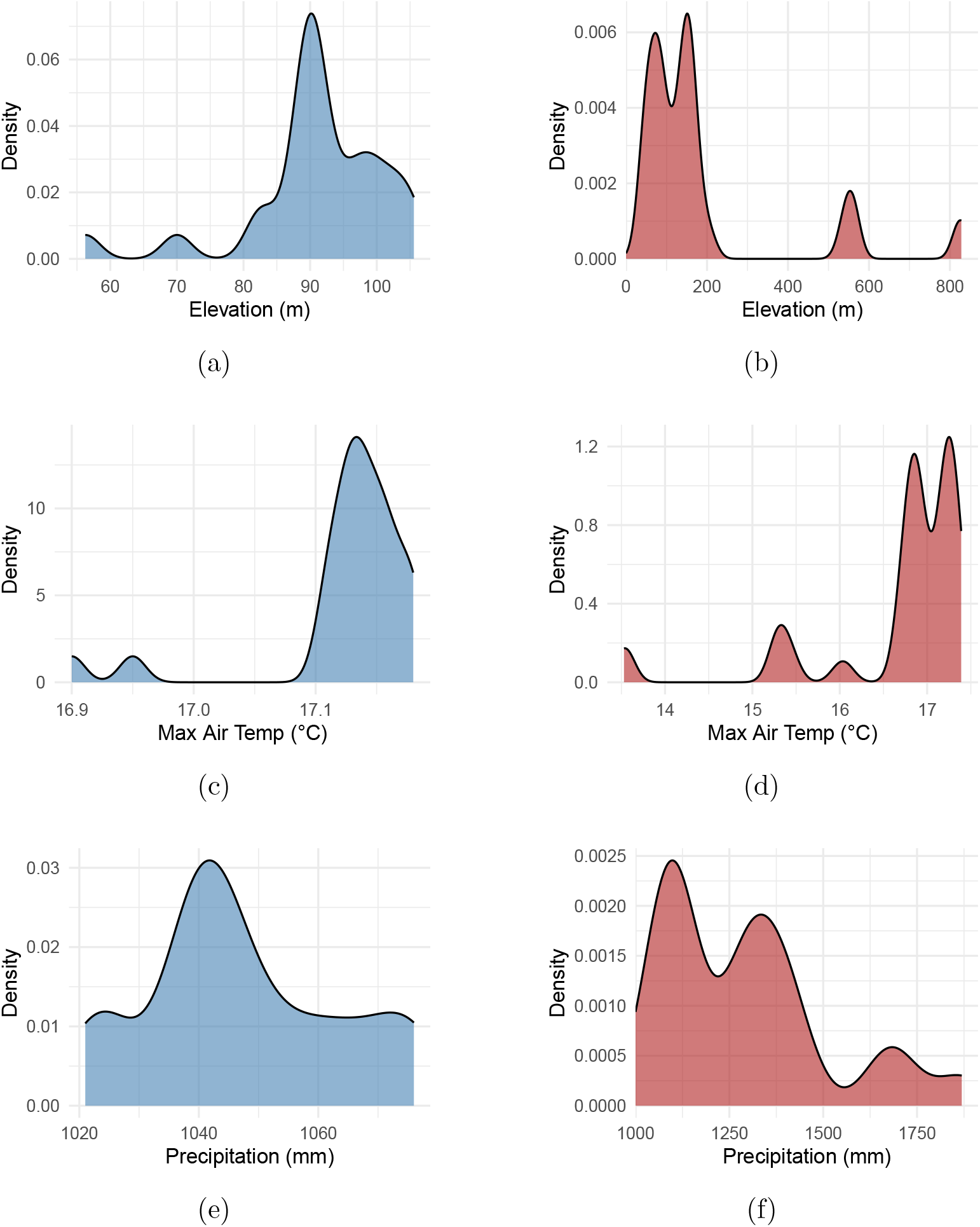
Comparison of the densities of elevation, temperature, and precipitation between Fanno Creek and the rest of the watersheds: Panels (a), (c) and (e) represent Fanno Creek while (b), (d), (f) represent the rest of the watersheds.

**Figure 9:**
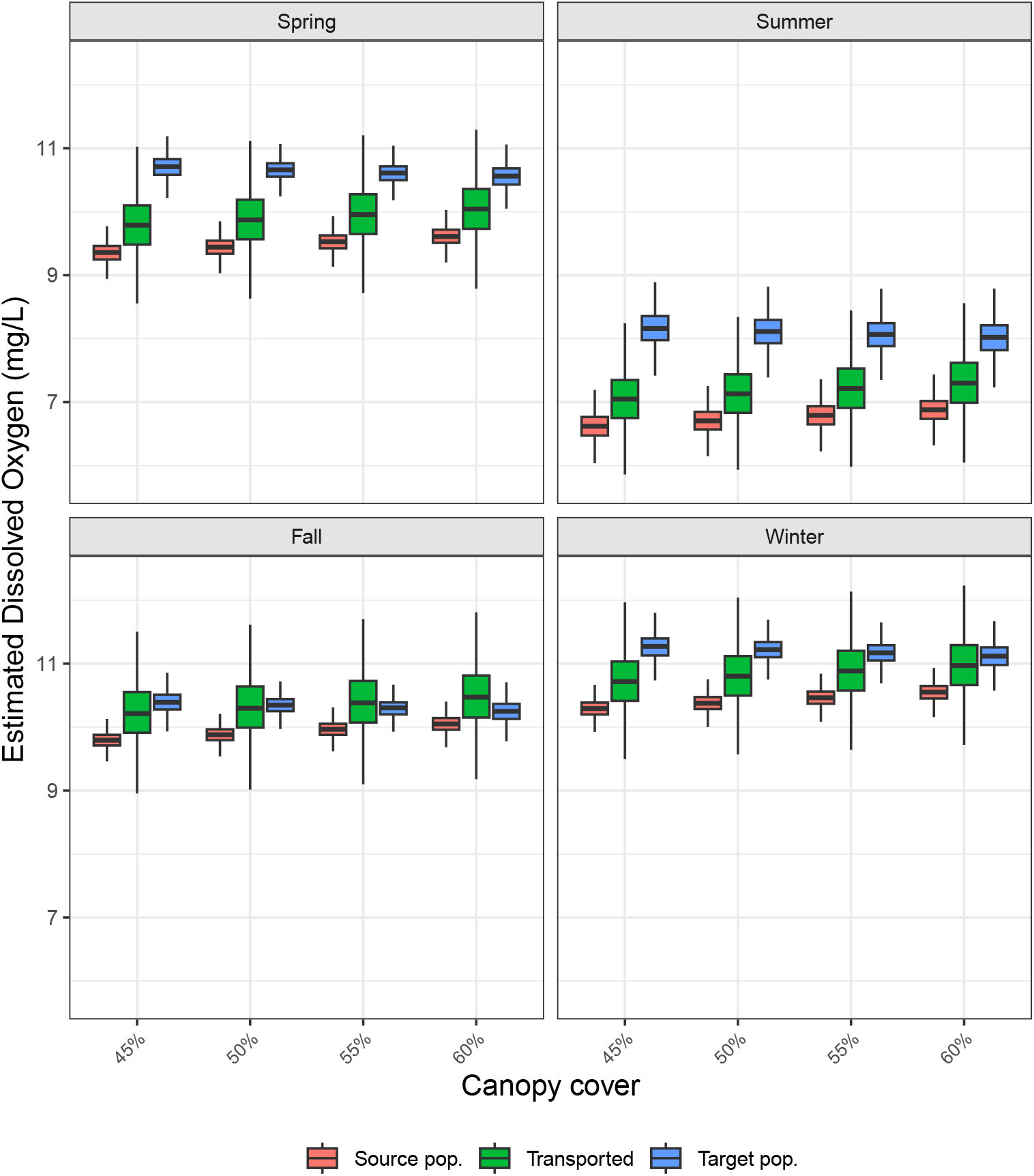
The results for the real-data case study for Johnson Creek (*N* = 115). The dissolved oxygen concentrations are reported separately for each season under the interventions do(*X* = 45%), do(*X* = 50%), do(*X* = 55%) and do(*X* = 60%), using three modeling approaches. Boxplots show median, interquartile range (IQR), and whiskers extending to 1.5 *×* IQR calculated from 2500 bootstrap iterations; outliers beyond the whiskers are omitted for clarity..

**Figure 10:**
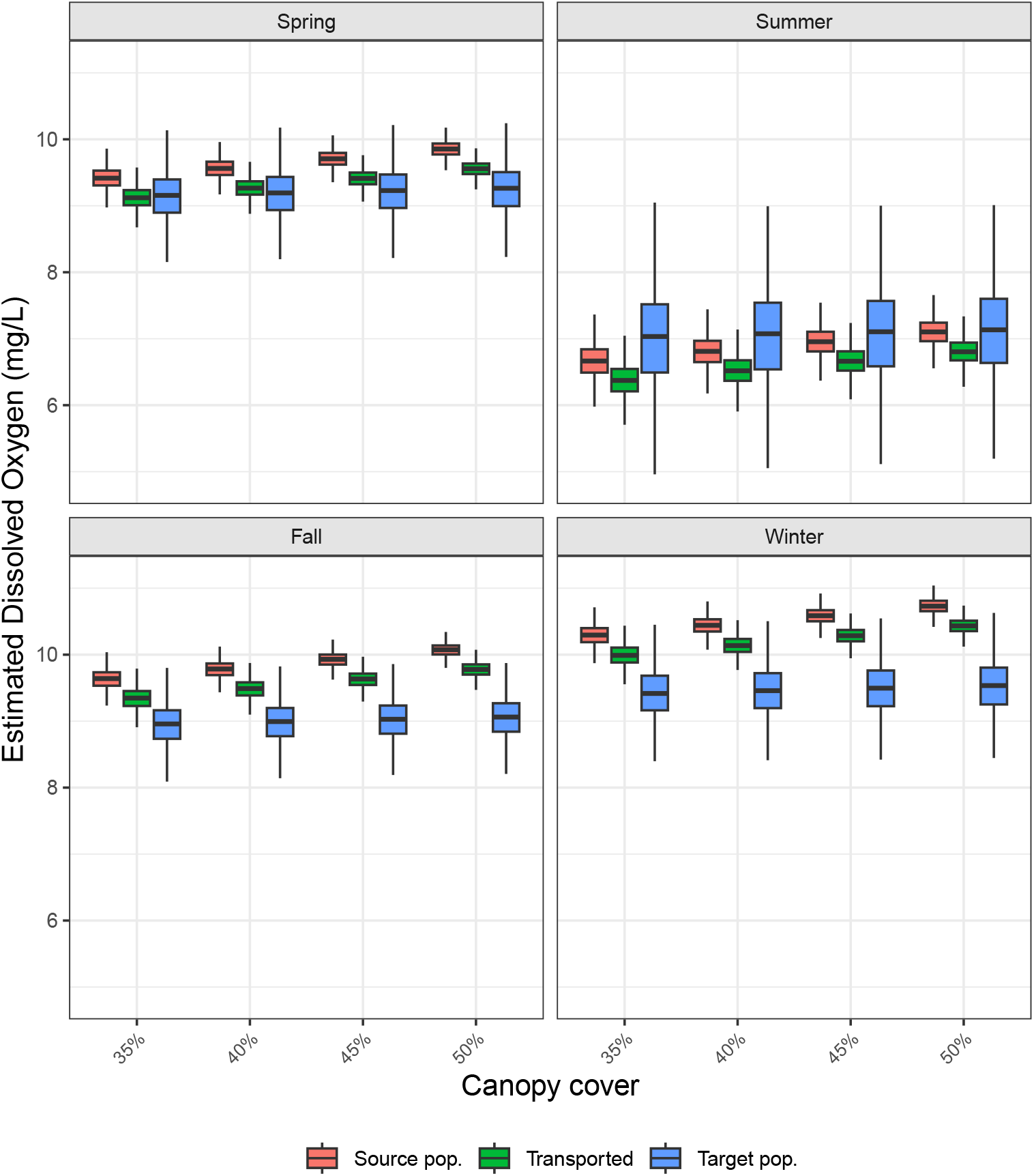
The results for the real-data case study for Kellogg Creek (*N* = 40). The dissolved oxygen concentrations are reported separately for each season under the interventions do(*X* = 45%), do(*X* = 50%), do(*X* = 55%) and do(*X* = 60%), using three modeling approaches. Boxplots show median, interquartile range (IQR), and whiskers extending to 1.5 *×* IQR calculated from 2500 bootstrap iterations; outliers beyond the whiskers are omitted for clarity.

